# BORD: A Biomedical Ontology based method for concept Recognition using Distant supervision: Application to Phenotypes and Diseases

**DOI:** 10.1101/2023.02.15.528695

**Authors:** Sumyyah Toonsi, Şenay Kafkas, Robert Hoehndorf

## Abstract

**Motivation:** Concept recognition in biomedical text is an important yet challenging task. The two main approaches to recognize concepts in text are dictionary-based approaches and supervised machine learning approaches. While dictionary-based approaches fail in recognising new concepts and variations of existing concepts, supervised methods require sufficiently large annotated datasets which are expensive to obtain. Methods based on distant supervision have been developed to use machine learning without large annotated corpora. However, for biomedical concept recognition, these approaches do not yet exploit the context in which a concept occurs in literature, and they do not make use of prior knowledge about dependencies between concepts.

**Results:** We developed BORD, a Biomedical Ontology-based method for concept Recognition using Distant supervision. BORD utilises context from corpora which are lexically annotated using labels and synonyms from the classes of a biomedical ontology for model training. Furthermore, BORD utilises the ontology hierarchy for normalising the recognised mentions to their concept identifiers. We show how our method improves the performance of state of the art methods for recognising disease and phenotype concepts in biomedical literature. Our method is generic, does not require manually annotated corpora, and is robust to identify mentions of ontology classes in text. Moreover, to the best of our knowledge, this is the first approach utilising the ontology hierarchy for concept recognition.

**Availability:** BORD is publicly available from https://github.com/bio-ontology-research-group/BORD

## 1 Introduction

Biomedical concept recognition is a language processing task that refers to extracting and normalising the mentions of biomedical concepts from text to structured resources such as ontologies. Automatic assignment of biomedical ontology concepts in unstructured text is a challenging task due to the use of ambiguous entities, abbreviations, synonymous entities, nested structures, and lexically variable description due to the use of natural language. Accurately recognising concepts in text facilitates the use of knowledge found in the literature in textual format for further analysis and various tasks. More specifically, linking literature and ontologies facilitates accessing, analysing and processing data efficiently. Furthermore, it enables performing downstream text mining tasks such as relationship extraction between the recognised entities, expansion of ontologies with the recognised new concept names and synonyms, as well as information retrieval for entities of interest.

Methods for concept recognition usually rely on lexical dictionary-based or supervised machine learning approaches. Lexical approaches such as the NCBO (National Center for Biomedical Ontology) annotator (Jonquet *et al.*, 2009), ZOOMA (Kapushesky *et al.*, 2011), and the OBO (Open Biological and Biomedical Ontologies) annotator (Taboada *et al.*, 2014) are not able to recognise new concepts and cannot detect all variations of expressions since their scope is limited to the lexical variations in the dictionaries. On the other hand, although supervised learning approaches have been applied on a wide range of biomedical concepts such as genes/proteins (Wei *et al.*, 2015) and diseases (Leaman *et al.*, 2013), they usually require large, manually annotated corpora which are not easily obtainable. The available labeled corpora are often insufficient to obtain a supervised model that can generalise to concepts uncovered by the labeled corpora. To address this, more recently, several distant-supervision based approaches have been proposed for concept recognition. In the distant supervision learning scheme, labels are learned based on a weakly labeled training set, i.e., obtained from an imprecise source (e.g., annotations generated by using rules or vocabularies). For example, PhenoTagger (Luo *et al*., 2021b) is a hybrid method that relies on a dictionary and a distantly supervised model trained only on the dictionary names, synonyms and their lemmas (the base form of a word found in a dictionary) to recognise concepts in text. Altogether, existing concept recognition methods utilise lexical signals, no or limited contextual information from relatively small labeled corpora. The contextual information found surrounding entities in text could also help guide the learning process and recognising concepts in text. However, existing methods leave the context in which concept are mentioned under-exploited. Furthermore, concepts are often structured hierarchically, in particular when the concepts are part of an ontology. However, existing concept recognition and normalisation methods rarely exploit this hierarchy to improve the concept recognition and normalisation process.

Here, we address two main research questions related to concept recognition and normalisation. The first question asks whether the context in which concepts are mentioned in text can be used without explicitly generating training corpora to develop a concept recognition method; for this purpose, we will exploit a distant supervision approach where training data is generated using lexical rules. The second question asks whether prior knowledge of hierarchical relations between concepts can be utilised to improve the performance of concept normalisation; to answer the second question, we develop an approach where information about super-concepts guides the concept normalisation, in particular when concepts are only partially matched.

We developed BORD as a generic concept recognition and normalisation method which does not require manually annotated corpora and is competitively able to identify mentions of concepts in text, in particular when the concepts are part of an ontology. BORD exploits both the lexical and contextual components of the concept mentions in text. To this end, BORD uses a dictionary constructed from the labels and synonyms of concepts to pre-annotate mentions of concepts in biomedical text. We then use these mentions in the text as weak (noisy) labels to train a language model (Lee *et al.*, 2020). To map the identified mentions in text to their concept identifiers, we developed a normalisation method that is inspired by the reciprocal best match approach but extended to incorporate hierarchical information. We applied BORD to the tasks of recognising phenotype and disease concepts based on the Human Phenotype Ontology (HPO) (Köhler *et al*., 2018) and the MEDIC vocabulary (Davis *et al*., 2012a). We evaluated BORD on two curated datasets covering diseases (Doğan *et al*., 2014) and phenotypes (Mohan and Li, 2019). Our results show that BORD outperforms state-of-the-art methods for recognising and normalising disease and phenotype concepts in text.

## 2 Materials and Methods

### 2.1 Ontologies and benchmark corpora

We generated and used two dictionaries to annotate abstracts from Medline (NCBI, 1996b). The first dictionary covers disease concepts and the second dictionary covers phenotype concepts. We used MEDIC, (downloaded on 1/March/2022) and the Disease Ontology (DO) (downloaded on 15/April/2022), for the disease concepts; DO is an ontology from the Open Biomedical Ontologies (OBO) (Schriml *et al.*, 2018), whereas MEDIC is a vocabulary of disease terms represented in the Web Ontology Language (OWL) (Davis *et al.*, 2012b). We used the Human Phenotype Ontology (HPO) (Köhler *et al.*, 2018) (downloaded on 5/Jan/2022) to generate the phenotype dictionary. For each of the dictionaries, we obtained the name and synonyms of each concept and further included the plural form of each entry (see Section 2.3 for further details).

To evaluate BORD, we used two benchmark corpora; the NCBI–Disease Corpus (Doğan *et al.*, 2014) and the MedMentions Corpus (Mohan and Li, 2019). We used the version of the NCBI–disease Corpus (Luo *et al.*, 2021a) released by Luo *et al.* (Luo *et al.*, 2021b) where the concepts are mapped to the Medical Subject Headings (MESH) (NCBI, 1996a) or the Online Mendelian Inheritance in Man (OMIM) catalog (Amberger *et al.*, 2014).

MedMentions is a large corpus annotated by an extensive set of Unified Medical Language System (UMLS) concepts. We selected the abstracts with phenotype annotations from MedMentions and named this the MedMentions–phenotype Corpus. We used UMLS-to-HPO mappings from UMLS (14,708 distinct HPO concepts are mapped to at least one UMLS concept) to obtain the HPO codes of the phenotype annotations. Table 1 shows the distribution of the abstracts and annotations in the two benchmark corpora.

**Table 1.** Distribution of abstracts and annotations in the benchmark corpora

| Corpus | Abstracts | Annotations |
| --- | --- | --- |
| NCBI–disease training | 593 | 5146 |
| NCBI–disease development | 100 | 788 |
| NCBI–disease test | 100 | 960 |
| MedMentions–phenotype training | 1291 | 6772 |
| MedMentions–phenotype development | 428 | 2287 |
| MedMentions–phenotype test | 405 | 2190 |
| MedMentions–disease test | 879 | 3726 |

Phenotypes are mapped to HPO; Diseases are mapped to MESH or OMIM

We used Medline (NCBI, 1996b) as a literature resource to train our models. To select abstracts, we used an in-house index covering 32,923,095 Medline records (downloaded on Dec-15-2021) generated using Elasticsearch (Elastic and Swiftype, 2010) for abstracts and titles (Uludag, 2021).

### 2.2 BORD system overview

The BORD system, depicted in Figure 1, consists of two phases; the training phase and the prediction phase. An ontology contains a controlled vocabulary expressed in an ontology representation language (Bodenreider, 2008). In the training phase, we first extract the vocabulary; more specifically all concept labels (names) and synonyms from a given ontology (MEDIC, DO, and HPO in our case) to form our initial dictionary. We then expand the vocabulary by generating the plural form of each term. Second, we use the dictionary to lexically annotate the literature creating a weakly annotated dataset. We then use the dataset to train a deep learning model (BioBERT (Lee *et al.*, 2019)). In the test (prediction) phase, we use the BORD model to identify the ontology concepts from the biomedical texts. Next, we map the identified mentions to their corresponding identifier in the ontology using an ontology-based normalisation approach.

**Fig. 1.**
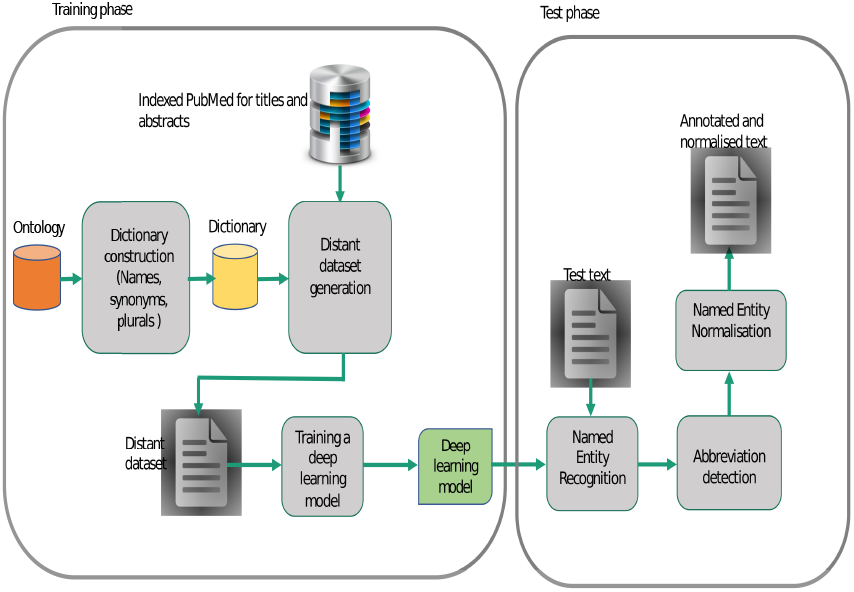
BORD System overview

### 2.3 Dictionary generation

To generate our dictionary, first, we downloaded the ontologies and extracted the names and synonyms of all concepts. Second, we filtered out the possible ambiguous names which are often stop words, short names (1 or 2 character long) and names shared by two different concepts from the dictionary. Filtering out ambiguous names is a common practice used in text mining workflows which rely on lexical matches. We used the Natural Language Toolkit (NLTK) stop words (Brigadir, 2019) and filtered out any exact match with the names/synonyms in MEDIC, DO and HPO. In MEDIC, DO, and HPO, we did not find any match with the list of stop words. We also filtered out the names/synonyms having less than 3 characters to avoid false positives. Additionally, for the generation of the dictionary for diseases, we filtered out all the disease names which exactly match with protein names/synonyms from the HUGO Gene Nomenclature Committee (HGNC) Database (Tweedie *et al.*, 2020). Third, we generated the plural form of each name/synonym by using the Inflect Python module (Dyson, 2022). For example, the module generates “malformations of lip” (HP:0000159) for the given multi-word term, “malformation of lip”. Our final disease dictionary covers 244,903 disease names and synonyms of 29,374 distinct concepts from MEDIC and DO. The final phenotype dictionary covers 79,010 phenotype names and synonyms of 14,631 distinct concepts from HPO.

### 2.4 Training dataset construction

To generate the training set for distant supervision, first, we retrieved the relevant literature by searching the indexed Medline for the exact match of each name/synonym from the dictionaries. We retrieved the top 5 Medline abstracts/titles hits per concept that is identified based on the default Elastic Search Engine relevance scoring settings (TF-IDF based scoring). Second, we used the dictionaries and annotated the downloaded abstracts lexically and converted the annotations to the I-O-B format (a common format for tagging tokens in a chunking task) (Ramshaw and Marcus, 1995) by using spaCy (Honnibal and Montani, 2017). Finally, we obtained two corpora; one for the disease concepts and the other for the phenotype concepts. We found 16,307 distinct phenotype names/synonyms belonging to 6,962 classes from HPO in at least one Medline record by searching the indexed literature. These concepts are covered by 74,087 distinct Medline abstracts/titles, and we used them as our training set for phenotypes. We found 35,333 distinct disease names/synonyms linked to 8,400 distinct concepts from MEDIC and DO in at least one Medline records. These concepts are covered by 187,462 distinct Medline abstracts/titles that we used as our training set for disease concepts.

### 2.5 Concept Recognition

We addressed the concept recognition task with two subtasks; Named Entity Recognition (NER) and Named Entity Normalisation (NEN). NER refers to identifying borders of entity mentions in text (e.g., disease and phenotype mentions). NEN refers to linking identified entities in text to the concepts defined in ontologies or databases. Figure 2 depicts the phenotype NER and NEN tasks on a sample sentence from the MedMentions–phenotype test dataset. NER is concerned with finding the words highlighted in red (“bone loss”, fractures, “oxidative stress”), while NEN is used to identify the HP identifiers pointed by the arrows.

**Fig. 2.**
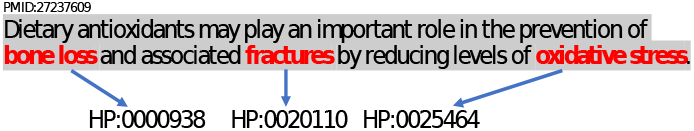
Sample phenotype concept recognition

#### 2.5.1 Named entity recognition using distant supervision

We used distant supervision to train a model by using BioBERT to recognise disease and phenotype mentions in text. BioBERT is a domain-specific language model; a BERT (Devlin *et al.*, 2019) pre-trained model based on large-scale biomedical corpora. BERT (Bidirectional Encoder Representations from Transformers) (Devlin *et al.*, 2019) is a contextualized word representation model trained using masked language modeling. It provides self-supervised deep bidirectional representations from unlabeled text by jointly conditioning on both left and right contexts. The pre-trained BERT model can be fine-tuned with an additional output layer to generate models for various desired NLP tasks. BERT has been widely used in Natural Language Processing and text mining. We used *simpletransformers* (Rajapakse, 2019) which provides a wrapper model to distantly supervise BORD’s entity recognition component. More specifically, the wrapped model is used to fine-tune BERT models by adding a token-level classifier on top that classifies tokens into one of the output classes which are I-O-B (Inside-Outside-Beginning). In the training phase, our model is initialised with weights from BioBERT-Base v1.1 (https://github.com/dmis-lab/biobert) and then fine-tuned on the disease and phenotype concept recognition task using our training corpora.

#### 2.5.2 Unsupervised entity normalisation

We normalised the tagged mentions of concepts by developing and using a method inspired by the Reciprocal Best Match (RBM) algorithm (Ward and Moreno-Hagelsieb, 2014). To this end, we used the dictionaries constructed according to 2.3, expanded them with lemmas, and further tokenised each name and synonym in the dictionaries to allow flexible matches. We subsequently matched the mentions obtained from the NER part to the names and synonyms in the dictionaries. We retrieved concept IDs for any concept that matches at least one token with the NER mentions and considered them as candidates. We assigned these candidates scores according to our RBM algorithm with the edit distance acting as the similarity measure as shown in Algorithm 1. Our aim is to find the best match for each mention token in the candidate tokens, if possible. The edit distance threshold dictates whether two tokens should be matched according to their lexical similarity.

#### Algorithm 1 RBM

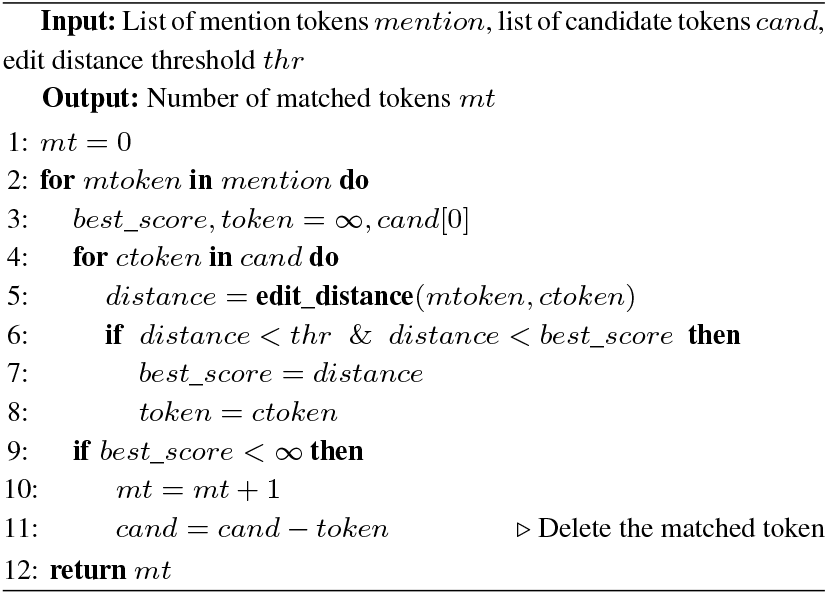

After matching tokens, we scored candidates based on two measures: Mention Matching Ratio (MMR), and Candidate Matching Ratio (CMR). MMR is the ratio of matched tokens between the candidate and the mention over the total number of tokens in the mention.

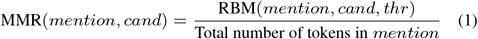

CMR is the ratio of matched tokens over the total number of tokens in the candidate.

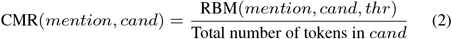

We obtained threshold values for the best performing MMR, CMR and edit distance for normalisation based on a grid search on the lexically annotated development sets. We varied both MMR and CMR [0.5,1] with a step of 0.1. Similarly, we varied the edit distance difference between [0.5,0] with a step of 0.1. The edit distance here represents how dissimilar we allow the matched tokens to be; the less, the more strictly similar they need to be. For ontology-based normalisation of MedMentions-phenotypes, we found that the best performing threshold values were: MMR=0.5, CMR=0.8, edit distance=0.1. For ontology-based normalisation of NCBI-diseases, we found that the best performing threshold values were: MMR=0.8, CMR=0.8, edit distance=0.1. It is important to note that beyond 0.5 thresholds, our normalisation performance is highly stable with multiple settings yielding the same results. Essentially, these two measures allow us to determine whether the number of matched tokens between the mention and the candidate is sufficient to predict the candidate. We declared the candidate of maximum score that passes these thresholds as a predicted concept ID for their its mention, if any.

Figure 4 depicts our normalisation method on a sample. First, we tokenise the mention “Hashimoto thyroiditis” into “Hashimoto” and “thyroiditis” then look up each token in the dictionary. Second, we retrieve any concept that matches at least one token from the mention as a candidate. We find two candidates: “Hashimoto thyroiditis” and “thyroiditis”. Third, we score these candidates such that each token from the mention is matched to a token in a given candidate. In the example, the “Hashimoto thyroiditis” candidate has a MMR of 2/2 because two of the mention tokens matched to the two tokens in the candidate. Similarly, it has a CMR of 2/2 because the two tokens of the candidates were successfully matched to the mention tokens.

**Fig. 3.**
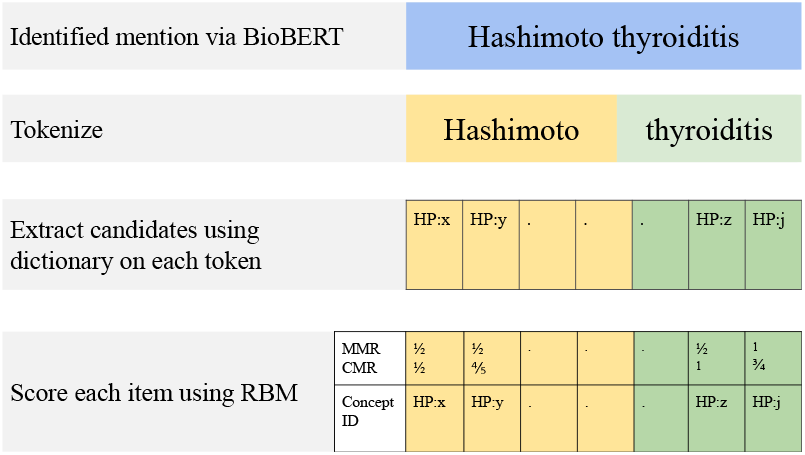
Normalisation method

**Fig. 4.**
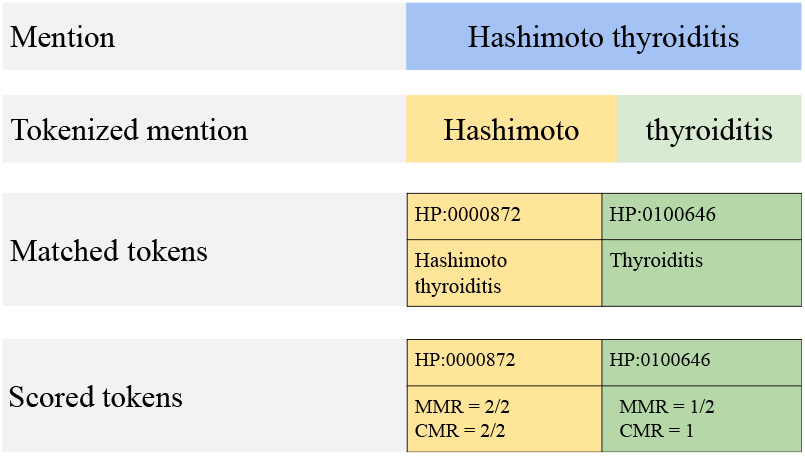
Example of normalisation

##### Using ontology hierarchy for entity normalisation

Due to the NER model recognizing entities partially in a subset of cases, some important information that guide the identification of the most specific class might be missing. Moreover, sometimes information that support a more specific class are not directly part of the mention but are rather mentioned somewhere else in the abstract. These two cases make mapping mentions to their direct concepts in ontologies challenging. We address these issues by exploiting the ontology structure in the mapping process as shown in Algorithm 2. For this purpose, if no candidate concept meets the thresholds for the MMR and CMR scores, we retain classes that have a CMR of 1. These are classes that exactly match only a part of the mention. We call such classes as general classes. For each such general class, we keep track of the tokens which were not matched with it, i.e. remaining tokens. We then consider the children of the general classes as candidates to match with the remaining tokens. If no children meet the MMR and CMR thresholds, we predict the general classes instead.

#### Algorithm 2 BORD normalisation

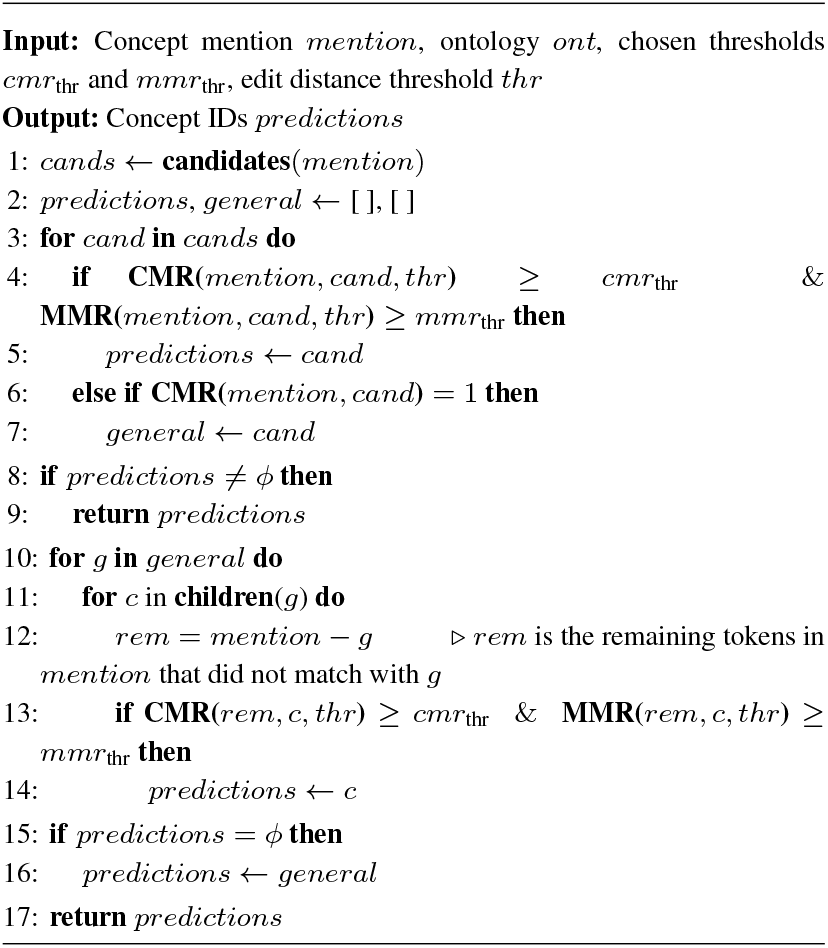

For example, the mention “neuroendocrine carcinoma of the breast” captured by the NER part, does not pass the MMR and CMR thresholds for any candidate. However, it matches the parent class HP:0003002 “Breast carcinoma”. Although we cannot find any child candidate that matches the remaining tokens “neuroendocrine”, we predict the parent class to be of coarser granularity.

## 3 Results

We applied BORD on three separate corpora covering phenotype and disease concepts; MedMentions–phenotype, MedMentions–disease, and NCBI–disease. We reported our NER and NEN results using the Precision, Recall and F-score metrics. We compared BORD’s performance against two state-of-the-art methods: Supervised model (BioBERT for NER and our normalisation for NEN) and distantly-supervised PhenoTagger. To compare against PhenoTagger, we trained the PhenoTagger disease model using their GitHub recommendations as no trained disease model was available. We used two types of evaluation: strict and relaxed. In the strict scheme, we only consider as true positives predictions that match the curated annotation boundaries perfectly, i.e., predictions having the same starting and ending indices as the curated annotations. In the relaxed scheme, we consider any partial overlap between the prediction and the curated annotations to be a true positive, i.e., they are positives whenever the indices of the prediction and the curated annotations overlap.

### 3.1 Context enhances NER performance

We examined BORD and several state-of-the art methods in NER to answer the first research question regarding the potential of context to improve the performance. These methods are: supervised BioBERT trained using small curated datasets, PhenoTagger which is trained by using labels and synonyms without context, and our unsupervised dictionary. Table 2 presents the performance of BORD and the aforementioned state-of-the-art methods in phenotype NER on the MedMentions–phenotype test dataset. With the inclusion of context at a large scale, BORD achieved the highest F1-score compared with all other methods.

**Table 2.** NER Performance on phenotype concepts (MedMentions–phenotype test set)

| Method | Strict |  |  | Relaxed |  |  |
| --- | --- | --- | --- | --- | --- | --- |
|  | P | R | F1 | P | R | F1 |
| BORD | 0.595 | **0.736** | **0.658** | 0.649 | 0.802 | **0.717** |
| BORD (Weak MM) | **0.596** | 0.589 | 0.593 | **0.663** | 0.655 | 0.659 |
| Supervised BioBERT | 0.570 | 0.734 | 0.642 | 0.613 | 0.788 | 0.690 |
| Dictionary | 0.446 | 0.676 | 0.540 | 0.510 | 0.757 | 0.609 |
| PhenoTagger | 0.480 | 0.716 | 0.575 | 0.544 | **0.809** | 0.650 |

We investigated if the use of context can help in recognising other concepts. To that end, we reported our model’s performance in detecting disease concepts on the NCBI–disease and MedMentions–disease test datasets in Table 3. Our results showed that supervised BioBERT performed the best on NCBI–disease because concepts are highly conserved in this dataset. To fairly compare the performance of methods, we carried out further evaluation on the MedMentions–disease dataset. BORD achieved the highest F1-score on MedMentions–disease showing its advantage of using context at a large scale.

**Table 3.** NER Performance on disease concepts

| Method | Strict |  |  | Relaxed |  |  |
| --- | --- | --- | --- | --- | --- | --- |
|  | P | R | F1 | P | R | F1 |
| **NCBI–disease** |  |  |  |  |  |  |
| BORD | 0.791 | 0.582 | 0.671 | 0.951 | 0.698 | 0.805 |
| BORD (Weak NCBI) | 0.819 | 0.570 | 0.672 | **0.972** | 0.675 | 0.797 |
| Supervised BioBERT | **0.839** | **0.886** | **0.862** | 0.916 | **0.958** | **0.937** |
| Dictionary | 0.526 | 0.519 | 0.523 | 0.788 | 0.742 | 0.765 |
| PhenoTagger | 0.676 | 0.606 | 0.639 | 0.883 | 0.770 | 0.822 |
| **MedMentions–disease** |  |  |  |  |  |  |
| BORD | 0.608 | 0.689 | **0.646** | 0.652 | 0.740 | **0.693** |
| BORD (Weak NCBI) | **0.657** | 0.558 | 0.604 | **0.723** | 0.616 | 0.665 |
| Supervised BioBERT | 0.510 | 0.669 | 0.579 | 0.580 | 0.770 | 0.66 |
| Dictionary | 0.473 | 0.638 | 0.543 | 0.541 | 0.723 | 0.619 |
| PhenoTagger | 0.502 | **0.718** | 0.591 | 0.566 | **0.803** | 0.664 |

To investigate whether the amount of context affects the performance, we observed the effect of varying the training set size of BORD. To this end, we weakly labeled the training sets of NCBI and MedMentions (smaller size with fewer contexts) and used them to train BORD (Weak MedMentions/NCBI). Results showed that a version of BORD that is trained on a larger amount of context outperformed BORD on less context (Weak MedMentions/NCBI) (Tables 2 and 3).

### 3.2 Ontology enhances NEN performance

To answer the second research question of whether the ontology can help guide the normalisation of mentions to ontology concepts, we compared BORD which uses ontology-based normalisation (see Tables 4 and 5) against other state-of-the-art methods. These methods include PhenoTagger and the dictionaries that we created. PhenoTagger maps mentions to ontology concepts through its model which is distantly supervised directly on the ontology class labels and synonyms. The dictionaries map mentions to ontologies through exact match. Results show that BORD, which uses an unsupervised normalisation that utilises the ontology hierarchy, achieved the highest F1-score on the phenotypes (Table 4). On the disease concepts, BORD or BORD (weak NCBI) outperformed the other methods in terms of F1-score (Table 5), demonstrating that our method improves concept normalisation.

**Table 4.** NEN Performance on phenotype concepts (MedMentions–phenotype test set)

| Method | Strict |  |  | Relaxed |  |  |
| --- | --- | --- | --- | --- | --- | --- |
|  | P | R | F1 | P | R | F1 |
| BORD | 0.587 | **0.726** | **0.649** | **0.621** | **0.769** | **0.687** |
| BORD (Weak MM) | **0.592** | 0.585 | 0.588 | 0.620 | 0.613 | 0.616 |
| Dictionary | 0.413 | 0.485 | 0.444 | 0.531 | 0.616 | 0.570 |
| PhenoTagger | 0.472 | 0.704 | 0.565 | 0.504 | 0.753 | 0.604 |

**Table 5.** NEN Performance on disease concepts

| Method | Strict |  |  | Relaxed |  |  |
| --- | --- | --- | --- | --- | --- | --- |
|  | P | R | F1 | P | R | F1 |
| **NCBI-disease** |  |  |  |  |  |  |
| BORD | 0.734 | 0.541 | 0.623 | 0.802 | 0.591 | **0.680** |
| BORD (Weak NCBI) | **0.768** | 0.534 | **0.630** | **0.828** | 0.576 | 0.679 |
| Dictionary | 0.482 | 0.485 | 0.484 | 0.5472 | 0.550 | 0.548 |
| PhenoTagger | 0.631 | **0.566** | 0.596 | 0.719 | **0.641** | 0.678 |
| **MedMentions-disease** |  |  |  |  |  |  |
| BORD | 0.588 | 0.666 | **0.624** | 0.608 | 0.689 | **0.646** |
| BORD (Weak NCBI) | **0.638** | 0.542 | 0.586 | **0.666** | 0.566 | 0.612 |
| Dictionary | 0.431 | 0.587 | 0.497 | 0.451 | 0.615 | 0.520 |
| PhenoTagger | 0.484 | **0.693** | 0.570 | 0.515 | **0.736** | 0.606 |

The discrepancy in the performances between BORD and BORD (weak NCBI) is more evident on the MedMentions–disease dataset than on the NCBI–disease data. This observation can be explained due to the highly conserved mentions in the test and training splits of the NCBI–disease dataset which helps to achieve competitive performance in NER even with a small amount of context usage (BORD (weak NCBI)).

### 3.3 Error Analysis

We manually analysed errors introduced by our ontology-based NEN and NER methods to gain more insights on the method that we developed. For this purpose, we randomly selected 20 False Positive (FP) and 20 False Negative (FN) annotations for the NER and NEN tasks separately from the NCBI–disease and MedMentions–phenotype test datasets. Table 6 shows the results from our analysis on the FP samples.

**Table 6.**
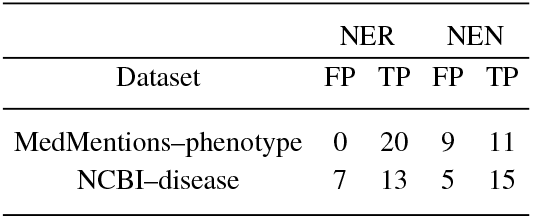
Error Analysis on False Positives

| Dataset | NER |  | NEN |  |
| --- | --- | --- | --- | --- |
|  | FP | TP | FP | TP |
| MedMentions-phenotype | 0 | 20 | 9 | 11 |
| NCBI-disease | 7 | 13 | 5 | 15 |

In the NER results, we found that all of the phenotype (20) and 13 out of 20 disease annotations that were FPs were actually True Positives (TPs), but they were not annotated in the curated datasets. For example, our model annotated the phenotype “high blood glucose” (HP:0003074) in “Diabetic neuropathic pain and high blood glucose were exhibited simultaneously …” (PMID:27461472) which is not annotated in the MedMentions–phenotype dataset. We found that the 7 FP disease annotations were due to ambiguous disease and gene abbreviations. For example, BORD annotated the abbreviation of “Wilms tumor 1” which is “WT1” in “Products of steroidogenic factor 1 (SF-1) and Wilms tumor 1 (WT1) genes are essential for mammalian gonadogenesis prior to sexual differentiation.” (PMID:9590178) as a disease name (Wilms tumor 1, MESH ID:D009396) where “WT1” is a gene in this specific context.

In the NEN results, we found that 11 phenotype and 15 disease mentions out of 20 FP annotations were actually TPs. These TPs were due to the cases where BORD normalises the mention to their corresponding classes in our dictionary by exact match but the mentions are mapped to different broader/parent) concepts in the annotated datasets. This is due to the used resources (e.g., MEDIC) having lexically identical synonyms for different concepts. For example, “bipolar effective disorder” in “Bipolar affective disorder (BPAD; manic-depressive illness) is characterized by episodes of mania and/or hypomania interspersed with periods of depression.” (PMID:9861003) is mapped to “major affective disorder 2” (MESH:C564108) by our method because it has “bipolar affective disorder” as an exact synonym. It is important to note that MEDIC can share the same exact synonym across multiple classes; there are eight classes (MESH:C567531, MESH:C567530, MESH:C567529, MESH:C567075, MESH:C567074, MESH:C567073, MESH:C565111, and MESH:C564108) sharing the “bipolar affective disorder” exact synonym. On the other hand, it is annotated with “bipolar disorder” (MESH:D001714) in the NCBI–disease Corpus. We found that the FPs (9 phenotypes, 5 diseases) in NEN were introduced mainly due to partial matches which can miss important tokens that are essential to find the correct concept identifer. For example, “colorectal adenomas and/or carcinoma” in “We have studied a set of 164 patients with multiple colorectal adenomas and/or carcinoma” (PMID:9724771) is mapped to “colorectal neoplasm” (MESH:D015179) in the NCBI–disease Gold dataset. However, our method maps it to a broader concept, “carcinoma” (MESH:D002277) since the NER model can capture only “carcinoma” from the whole mention, “colorectal adenomas and/or carcinoma”.

Table 7 shows the results from our analysis on the False Negative samples. We found that all the FNs in the NER task were missed since the BioBERT based NER model of BORD failed at capturing the term variations in the text. For example, our model did not recognise the phenotype “Reduces body Weight” in “A Novel Selective Inhibitor […] Reduces Body Weight in Diet -Induced Obese C57BL/6J Mice” (PMID:27832159). This specific phenotype is mapped to two alternative HPO classes which are “Decreased body weight” (HP:0004325) and”Weight loss” (HP:0001824) by the curators of MedMentions–phenotype Corpus. These two phenotype names from HPO are also the names that we used to distantly supervised our BioBERT-based NER model. Since these names are lexically different from the mention “Reduces body Weight”, our NER model misses it.

**Table 7.**
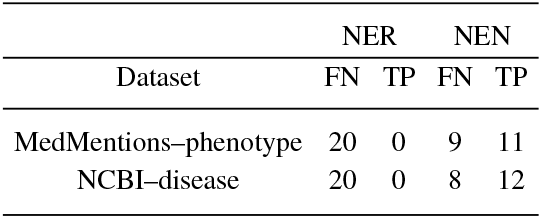
Error Analysis on False Negatives

The FN disease and phenotype samples that we analysed in the NEN task should be also considered as FPs. Because our method captures those annotations at the NER level correctly but maps them to different classes compared to the curated labels. Hence, a given annotation is treated as an FN (missed) according to the curated dataset but it is also an FP since our method maps the mention to another class (compared to the curated dataset) in the normalisation process.

### 3.4 Ablation studies

We further demonstrated the contribution of each component of our method on the MedMentions–phenotype and the NCBI–disease datasets. Because BORD can map mentions to broader classes in the ontology, we consider a special type of evaluation named “ont-evaluation” in Table 8. In this evaluation, we expanded the curated sets so that they include the direct parents of the assigned classes. For instance, we allow “Alzheimer Disease” (MESH:D000544) and/or “Dementia” (MESH: D003704) to be correct mappings to any mention of Alzheimer Disease.

**Table 8.** Ablation

| Component | Strict |  |  | Relaxed |  |  |
| --- | --- | --- | --- | --- | --- | --- |
|  | P | R | F1 | P | R | F1 |
| **MedMentions–phenotype** |  |  |  |  |  |  |
| Raw NER | 0.575 | 0.744 | 0.648 | 0.629 | 0.813 | 0.709 |
| Filtered NER | 0.595 | 0.736 | 0.658 | 0.649 | 0.802 | 0.717 |
| NEN | 0.587 | 0.726 | 0.649 | 0.621 | 0.769 | 0.687 |
| NEN ont-evaluation | 0.590 | 0.730 | 0.652 | 0.639 | 0.791 | 0.707 |
| **NCBI–disease** |  |  |  |  |  |  |
| Raw NER | 0.738 | 0.66 | 0.697 | 0.909 | 0.808 | 0.856 |
| Filtered NER | 0.791 | 0.582 | 0.671 | 0.951 | 0.698 | 0.805 |
| NEN | 0.734 | 0.541 | 0.623 | 0.802 | 0.591 | 0.680 |
| NEN ont-evaluation | 0.740 | 0.545 | 0.627 | 0.871 | 0.640 | 0.738 |

Filtering NER based on mappable candidates yielded a higher F1-score than the raw predictions. Moreover, when we considered the special “ont-evaluation”, we observed better normalisation performance. Namely, the relaxed evaluation improved by 2% on MedMentions–phenotype. On NCBI–disease the improvement is more evident as the overall F1-score increased by 5%. These results suggest that our ontology-based normalisation helps in resolving partial matches.

## 4 Discussion

Our main goal was to develop a method that can recognise concepts based on a given ontology or vocabulary without the requirement of a manually annotated corpora. To that end, we exploited textual context information and ontology structure for concept recognition. We observed that utilising the context via distant supervision improves NER performance. Although this was evident in our phenotype recognition results (see Table 2), it was not demonstrated on the NCBI–disease dataset. Investigation on these datasets revealed that there is a high overlap (around 80%) between the concepts in the training and testing sets. Hence, the model which was trained on a restricted set of mentions (BORD (Weak NCBI)) achieved similar performance as BORD in terms of F-score. However, when we applied our method on the MedMentions–disease dataset, BORD outperformed BORD (Weak NCBI). In addition, BORD which was trained on the lexically annotated Medline abstracts consistently had higher recall showing that it is more general and flexible to term variations. Another lesson learned by switching the dataset for diseases from NCBI to MedMentions is that BORD is more robust to changes in datasets and can generalize better compared to PhenoTagger.

We also observed that including the ontology structure as background knowledge provides a way to map partially matched mentions to close classes in the ontology. In particular, partially captured mentions can miss critical parts that guide the identification of the most specific concept. In this case, our method normalises to a concept of a slightly coarser granularity. This was evident in the ablation studies (NEN ont-evaluation) when the direct parents were considered as true positives during evaluation. Especially on the NCBI–disease dataset, we observed a substantial discrepancy between the regular evaluation (NEN) and ontology-based evaluation (NEN ont-evaluation). The MEDIC vocabulary was mainly built by integrating OMIM terms into the MeSH hierarchy (Davis *et al.*, 2012a), which provides a vocabulary that is less structured than a standard ontology.

Overall, while BORD outperformed two state-of-the-art methods (supervised and distantly supervised) on the task of recognising phenotype concepts, it achieved mixed results on disease concepts. This can be explained by nature of the NCBI–disease dataset which has high overlap between its test and training datasets. Furthermore, our manual error analysis revealed that some false positives should in fact be true positives in the curated datasets. This can be attributed to human curators missing annotations due to unclear boundaries between disease and phenotype concepts. We also found cases where the curators mapped concepts to broader classes even when information for a more specific class was available.

We identified several limitations in our work. BORD cannot identify nested mentions as the trained BERT-based NER model operates on a dataset represented in the standard I-O-B format. In addition, BORD also shares BioBERT’s limitations in recognising name variations in text as we observed through manual error analysis. Lastly, although BORD’s normalisation is designed to be flexible to lexical variations, mentions that are lexically distant from the curated concepts cannot be normalised. We alleviated some of these cases through the incorporation of the ontology structure.

## 5 Conclusion

We developed BORD, a Biomedical Ontology based method for concept Recognition using Distant supervision. BORD is generic and can be applied on any concept given an ontology or a vocabulary. Its main advantage is its ability to utilise a weakly labeled dataset for training. This allows BORD to be used for concepts where no curated corpora are available. While the use of an ontology is highly recommended because BORD uses the ontology hierarchy for concept recognition, it can still be used given any vocabulary. BORD is freely available at https://github.com/bio-ontology-research-group/BORD.

## Acknowledgements

We thank Dr. Mahmut Uludağ for his technical assistance in processing MEDLINE data.

